# Sex and genetic specific effects on behavioral, but not metabolic, responses to a high fat diet in heterogeneous stock rats

**DOI:** 10.1101/2022.03.25.485743

**Authors:** Aaron W Deal, Andrew Thurman, Osborne Seshie, Alexandria Szalanczy, Angela Beeson, Mackenzie Cockerham, Ellen L Risemberg, Anne Lenzo, Noelle Ozimek, Carl Langefeld, William Valdar, Leah C Solberg Woods

**Affiliations:** Department of Internal Medicine, Wake Forest School of Medicine, Winston Salem, NC, 27101; Department of Internal Medicine, University of Iowa College of Medicine, Iowa City, IA, 52242; Department of Genetics, University of North Carolina at Chapel Hill, Chapel Hill, NC, 27599; Curriculum in Bioinformatics and Computational Biology, University of North Carolina at Chapel Hill, Chapel Hill, NC, 27599; Department of Biostatistical Sciences, Wake Forest School of Medicine, Winston Salem, NC, 27101

**Keywords:** Diet-induced obesity, Diet, Outbred, Heterogeneous Stock, Heritability, Gene by diet, Obesity, Depression, Anxiety

## Abstract

Obesity is a growing epidemic associated with a range of comorbidities, including anxiety and depression. Genetics and environmental factors such as diet contribute to both adiposity and anxiety/depression. Heterogeneous stock (HS) rats are an outbred colony and useful for genetic mapping of complex traits. We have previously shown that HS male rats exhibit worsened metabolic and behavioral health in response to high fat diet (HFD). This study aims to determine if females have similar response to diet and if response to diet interacts with genetic background. We measured multiple metabolic (body weight, fat pad weight, glucose tolerance, fasting glucose and insulin) and behavioral (elevated plus maze, open field test, and forced swim test) outcomes in a large cohort of male and female rats on either HFD or low fat diet (LFD). We estimated overall heritability as well as heritability of response to diet for each outcome. Both sexes showed worsened metabolic measures when fed HFD compared to LFD. In contrast, only males exhibited altered behavioral responses to HFD relative to LFD, with no effect in females. Most metabolic and behavioral measures showed overall heritability in both sexes. In contrast, although there was some evidence for gene by diet (GxD) interactions for behavioral measures in males, GxD interactions were generally not found for the metabolic measures. These data demonstrate an important role of diet, sex and genetics in metabolic and behavioral phenotypes in HS rats, with a potential role of gene by diet interactions for behavioral outcomes only in males.

## Introduction

In 2018, more than 42% of US adults were obese ^1^, and prevalence continues to rise^2^. In addition to negative metabolic effects, including cardiovascular disease and diabetes^3^, obesity is associated with affective disorders in humans, such as anxiety^4^ and depression^5^. Although the causal relationship between obesity and behavioral health is uncertain, a meta-analysis found a bi-directional association between obesity and depression in humans, indicating both diseases contribute to development of the other^5^. Despite a strong link between obesity and anxiety/depression, mechanisms underlying the association remain unclear.

Both obesity and affective disorders are complex traits, driven by genetics and the environment. There is evidence that shared genetics is one potential factor contributing to the association between obesity and anxiety/depression. Specifically, human genome wide association studies (GWAS) have identified more than 900 loci associated with BMI^6–8^ and more than 100 loci associated with depression^9^ with some shared factors having been identified^10^. In addition to shared genetics, sex is known to play a role in the prevalence of symptoms of depression^11^ and anxiety^12^, where rates are higher among women. Additionally, while rates of obesity were not significantly different between American males and females in 2018, women had a higher rate of severe obesity compared to men^1^.

In addition to genetic susceptibility, overconsumption of a high fat diet (HFD) is a common environmental cause of obesity in humans^13^. Animal models of HFD-induced obesity also show expected negative effects on metabolic measures^14–17^, as well as anxiety-like and/or despair-like phenotypes ^14,18–21^. To better understand the relationship between obesity and anxiety/depression, an animal model would be highly valuable. Due to the complexity of known and unknown genetic factors contributing to both obesity and anxiety/depression, a highly genetically diverse model is likely to provide insights into the phenotypic and genetic variability associated with each disease that may be not be available through use of inbred models.

The outbred heterogeneous stock (HS) rat population was created from 8 inbred strains^22^ and maintained in a way to maximize genetic diversity. The highly diverse genetic background within the HS population makes it a powerful tool for genetic fine-mapping to small genomic intervals^23^ and has previously been utilized to identify genetic loci and candidate genes for insulin resistance, glucose tolerance, adiposity, and multiple behavioral measures^24–28^. Previous work by our group has shown that both adiposity and behavioral traits are heritable in HS male rats^26,29^. We have also shown that male HS rats are susceptible to the negative effects of HFD on metabolic and behavioral outcomes^14^.

In the current study, we investigated the effect of HFD on metabolic and behavioral phenotypes in HS male and female rats. In addition to effect of diet on phenotypes, we estimated overall heritability of phenotypes, as well as heritability of response to diet. The overall goal of the current study is to determine whether genes interact with diet to influence metabolic and behavioral phenotypes, as well as establish an animal model to study the relationship between obesity and behavioral health. We hypothesize that both male and female HS rats will show worsened metabolic and behavioral phenotypes in response to HFD and heritability estimates of these data will show gene by diet interaction in both sexes.

## Methods

### Heterogeneous stock rats

The HS rat colony was established in 1984 by the NIH by breeding together 8 inbred founder strains: ACI/N, BN/SsN, BUF/N, F344/N, M520/N, MR/N, WKY/N, and WN/N, and maintaining the colony in a way that minimizes inbreeding^22^. The HS colony was maintained at the Medical College of Wisconsin since 2006 with a colony set up at Wake Forest School of Medicine (WFSM) in 2017^23^. The animals in the current study come from the WFSM colony which has been named NMcwiWFsm:HS (Rat Genome Database number 13673907). At the time of these studies, the colony was maintained using 64-90 breeder pairs in a way that takes into account family relationships using a kinship coefficient to ensure that only distantly related animals are mated. Rats were weaned at 3 weeks of age and placed 2 per cage (non-sibling cage mates) for the remainder of the study. To maximize genetic diversity, we used no more than 2 males and 2 females from each breeder pair, placing siblings on separate diets. Animals were housed on a 12h light/dark cycle (700-1900). All protocols were approved by the Institutional Animal Care and Use Committee at WFSM.

### Diet

High fat diet (HFD) and matched low fat control diet (LFD) was purchased from Research Diets (New Brunswick, NJ; high fat chow, 5.21 kcal/g, 60% fat, 20% carbohydrate, 20% protein, cat. D12492; low fat chow, 3.82 kcal/g, 10% fat, 70% carbohydrate, 20% protein cat. no D12450J).

### Study design

#### Previous work in HS male rats

Two separate studies were previously conducted to assess the effect of HFD on metabolic and behavioral phenotypes in HS male rats^14^. To increase power for heritability estimates and diet effects, we have combined data from these previous studies with the larger study described below. The methods and results of those studies have previously been described in full^14^. In brief, a metabolic study investigated HFD effects on body weight, visceral fat pad weight, fasting glucose and insulin, and glucose tolerance, as well as passive coping behavior via forced swim test. HS male rats were assigned to HFD (n = 48) or LFD (n = 24) at 6 weeks of age. Based on the results of the passive coping behavioral test, we initiated a second study in HS males to more extensively investigate the effects of HFD (n = 32) versus LFD (n = 32) on behavior, specifically anxiety-like (open field test and elevated plus maze) and despair-like/passive coping (forced swim test and splash test) behavior. In the behavioral study, rats started diet at 4 weeks of age. Time-line of testing for these previous studies differ slightly from the larger study described below and are shown in **Supplementary Figure 1.**

#### Expanded study to test the role of HFD on metabolic and behavioral outcome in male and female HS rats

Based on the previous studies in HS male rats^14^, we initiated an additional study to assess both behavioral and metabolic phenotypes in response to HFD in a large number of both male and female HS rats. In this study, 189 male HS rats were placed on HFD (n = 95) or LFD (n=94) and 188 female HS rats were placed on HFD (n = 94) or LFD (n = 94). Most rats started diet at 6 weeks of age, with a subset of males and females (HFD: n = 24, LFD: n = 24, per sex) starting diet at 4 weeks of age. Time on diet prior to behavioral testing was equivalent (8 weeks) regardless of age at diet start. The study time-line is described in **Table 1.** To increase sample size in males, data from this larger study were combined with data from the previously published work as described in the data analysis section below. Total number of males and females for each phenotype is given in **Supplementary Table 1.** Numbers differ across tests because not all phenotypes were completed in all the animals due to issues caused by Covid-19 laboratory shut-downs.

**Table 1.** Timeline of metabolic and behavioral tests.

| Diet Week | Procedure |
| --- | --- |
| 8 | EPM |
| 9 | OFT |
| 11 | FST |
| 12 | IPGTT |
| 13 (21, n=64) | Necropsy |

### Outcomes

All metabolic and behavioral tests utilized in this study have been previously described as performed by our lab^14^. Tests for anxiety-like behavior included elevated plus maze (EPM, 8wk on diet) and open field test (OFT, 9wk on diet), while we used the forced swim test (FST, 11wk on diet) to test for despair-like/passive coping behavior (see **Table** 1). Metabolic tests included an intraperitoneal glucose tolerance test (IPGTT) administered after 12wk on diet, at which point serum was collected for fasting insulin levels (ALPCO, Salem, NH, cat. no 80-INSRTU-E10), as well as fasting glucose, final body weight, and visceral fat pad weight after 13wk on diet.

#### EPM

Rats were placed at the junction of 2 elevated, perpendicular arms, one axis with walls and one axis without walls (axis length 111.8cm, wall height 74.0cm, floor to apparatus 45.7cm). The location of the rat was monitored and timed via beam break at arm entrances. Time spent in closed or open arms was measured over 5 minutes (Med-PC IV, Med Associates, Inc. Fairfax, VT). Anxiety-like behavior is associated with increased time in the closed arms^30,31^.

#### OFT

Rats were placed in an empty arena (40.8 x 40.8 x 30.1cm) and allowed to explore for 30 minutes. Behavior was monitored via infrared beam break and collected for analysis using Omnitech tracking software (Omnitech Electronics Inc. Columbus, OH). Data collected were analyzed for total distance, movement and rest episodes, rearing episodes and center time, both after 5 minutes of the test and again after 30 minutes. Anxiety-like behavior is associated with decreased time spent in the arena center and hyperlocomotion^32,33^.

#### FST

Rats were placed in a tank of water (25°C±2, diameter 28.8cm, height 49.8cm, water depth 39cm) for 15 minutes on day 1 and 5 minutes on day 2 as previously described^34^. Video recording of day 2 was used to manually score movements made by the rat at 5 second intervals: immobility (floating), swimming, climbing, and diving. Passive coping^35^ and/or depressive-like behavior is associated with increased immobility^20,21,34,36^.

#### Intraperitoneal Glucose Tolerance Test

IPGTT was run as previously described^14^. Briefly, rats were fasted for 16 hours (1730-930) prior to testing. Blood glucose levels were measured after fasting and rats were then given glucose at 1g/kg body weight via intraperitoneal injection. Blood glucose was measured at 15, 30, 60, and 90 minutes after glucose injection. Area under the curve (AUC) was determined by applying the following equation: t*(x+y)/2, where t is the elapsed time in minutes, x is the initial glucose concentration (mg/dL), and y is the final glucose concentration, to each time period (15, 30, 60, and 90min). We then summed the results to determine each rats’ IPGTT AUC.

#### Necropsy

Animals were fasted for 16 hours (1700 – 900) prior to necropsy. Euthanasia was performed via live decapitation. Final body weight and serum were collected, along with fasting glucose and multiple tissues, including visceral fat pads (retroperitoneal, gonadal, and omental).

### Data analysis

We omitted 200 data-points for fasting insulin because they fell outside the range of the standard curve; we omitted 32 data-points from EPM because of lighting differences between mazes that impacted rat movement. All phenotypes were transformed to closely reflect a normal distribution within each diet and sex prior to analysis. Transformations were chosen using the Box-Cox method^37^ and are listed in **Supplementary Table 2.** For males, we combined the previously collected data with the data^14^ from the expanded study. All data were pooled and analyzed for batch effects and age at diet start within and between studies via 2-way ANOVA. Any variables shown to have a significant effect were accounted for in the linear mixed model described below. To evaluate the effect of diet on behavioral tests (EPM, OFT, and FST) and metabolic outcomes (body weight, fat pad weights, glucose AUC, and fasting glucose and insulin), we used package Ime4^38^ in R^39^ to build linear mixed effects models of the form:

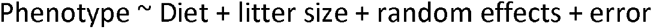

in which Diet and litter size (to account for wean weight) are fixed effects. Random effects included sibling status (to account for the complex family relationships of the HS), study e.g., (previously published data or the current expanded study), and within study cohort (to account for batch variation). Age at diet start did not impact any of the measures and was thus not included in the model. Data were analyzed separately for each sex. The results of the linear mixed effects model were corrected for multiple comparisons via Benjamini-Hochberg procedure (FDR = 0.05 or 0.1).

Phenotypes were analyzed for correlations between metabolic and behavioral outcomes within sexes overall and within sexes by diet. Pearson’s correlation coefficients were calculated and significant correlative relationships were determined following Benjamini-Hochberg correction for multiple comparisons (FDR = 0.05).

Factor analysis was utilized to probe for latent factors within the metabolic and behavioral responses to diet. The analysis was performed in R under the “psych” package^40^. Varimax rotation was used and the number of factors analyzed was determined via scree plot. To test for the effect of diet on the metabolic and behavioral data using fewer dimensions, a “factor phenotype” was calculated for each rat by summing phenotypic values weighted by the associated factor loading. The “factor phenotypes” were tested for significant effects of diet in the linear mixed effects models described above.

### Estimating Heritability and Gene-by-Diet Heritability

#### Statistical Model

For each combination of phenotype and sex, we estimated 1) the heritability, 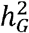, defined as the overall effect of genetics on the phenotype regardless of diet treatment, and 2) the gene-by-diet heritability 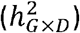, defined as the overall effect of genetics on the phenotypic response to diet treatment. Estimation was based on a linear mixed model that included fixed effects of litter size, study and diet, a random effect of cohort, and two polygenic effects, one for additive genetics and one for additive genetics interacted with diet. In more detail, for a given combination of sex and phenotype, *y_i_*, the phenotype of rat *i* ∈ {1,…, *n*}, was modeled as

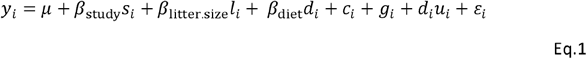

where *μ* is an intercept, *s_i_* indicates the study batch (1 or 2) and has effect *β*_study_, *l_i_* encodes the litter size (from 1 to 15) and has linear effect *β*_litter.size_, *d_i_* indicates diet treatment as *d_i_* = 1 for HFD and *d_i_* = −1 for LFD such that the overall effect of diet treatment is *β*_diet_; *c_i_* is a random intercept for cohort defined as 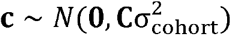, where 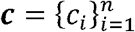 and **C** is an *n* × *n* kernel matrix with element {*C*}*_ij_* equal to 1 if rats *i* and *j* are in the same cohort and zero otherwise; *g_i_* and *u_i_* are polygenic effects modeled as 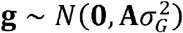 and 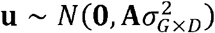 respectively, where A is the additive relationship matrix based on the pedigree (calculated using the R package “AGHmatrix”^41^), and 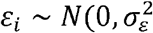 is independent, identically distributed residual noise.

The variance 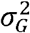 represents the contribution of additive genetics to a treatment-neutral version of the phenotype. Specifically, consider two genetically distinct rats that are otherwise identical in their study, cohort and litter, and where both receive a hypothetical, neutral treatment *d_i_* = 0. The difference in their expected phenotype (the phenotype minus residual noise) would be

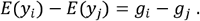

such that the magnitude of their difference is due to 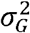.

The variance 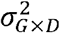 represents the contribution of additive genetics to individual-specific deviations from the population average diet response. Specifically, denote the phenotype of rat *i* under HFD as *y*_*i*,HFD_ and the phenotype of the same rat under the counterfactual scenario of it being given LFD as *y*_*i*,LFD_. The difference in the expected values of these two potential outcomes for rat *i* is

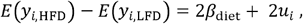

such that the extent of polygenic variation around the overall effect *β*_diet_ is due to 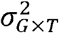. The above equation is analogous to that in models of genetic effects on the phenotypic difference between alternate-treated isogenic individuals in panels of inbred strains^42,43^, and our model is similar to the gene-by-environment interaction model proposed by Qu et al.^44^.

Heritabilities were defined as follows. First, define the intraclass correlation coefficient (ICC) of variance component *k* ∈ [*G, G × D*, cohort, *ε*} as proportion

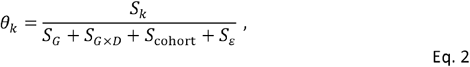

where 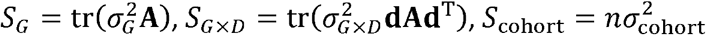 and 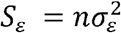. The heritability is then defined as 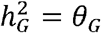 and the gene-by-diet heritability as 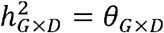.

#### Priors and Bayesian estimation

The linear mixed model in Eq 1 was estimated as a Bayesian model using Markov chain Monte Carlo (MCMC). The intercept and fixed effects were given improper flat priors, ie, *p*(*β*) ∝ 1. The random effect variances were given inverse gamma priors, ie, 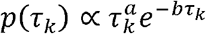, where 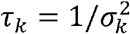 for variance component *k*, with shape *a* = −1 and rate *b* = 0. These hyperparameter values chosen so that the implied prior distribution *p*(*θ_k_*), and therefore the prior on 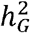 and 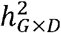, allows any value between 0 and 1 but is weighted towards lower values (between 0 and 0.4) and has moderate (but not infinite) weight at zero.

MCMC samples were generated using a Gibbs sampler written in R. (The conditional distributions used, which were relatively standard, are provided in the Appendix.) Posterior samples from *p*(*θ_k_|**y***) were generated as functionals of samples from 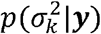 using Eq 2. For each combination of phenotype and sex, we ran the Gibbs sampler in 3 chains for 1 × 10^5^ iterations each. The raw samples from each chain were then thinned at 1:10 after discarding 1000 burn-in samples to give a final 29,700 samples from which we obtained point estimates as the posterior median and interval estimates as the highest posterior density (HPD).

#### Testing for non-zero heritability using Bayes factors

To quantify the evidence in favor of non-zero heritability or non-zero gene-by-diet heritability, we used Bayes factors^45^. The Bayes factor for a given *θ_k_* (eg, 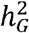 or 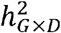) is defined as

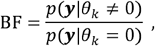

that is, the ratio of the marginal likelihood of the data when *θ_k_* is non-zero (ie, present) vs when it is zero (ie, absent). For example, in the case where *θ_k_* represents the gene-by-diet heritability, 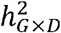, BF > 1 implies positive evidence for this heritability, BF < 1 implies evidence against it (ie, evidence for it being absent), and BF = 1 implies that the data provide no information either way. More generally, Kass and Raftery^45^ proposed the following guidelines for interpreting BF values (log_10_ BF in parentheses): <0.001 (less than −2) decisive evidence against, 0.001-0.1 (−2 to −1) strong evidence against, 0.1-0.3 (−1 to −0.5) substantial evidence against, 0.3-3.2 (−0.5 to 0.5) not worth mentioning (ie, inconclusive), 3.2-10 (0.5-1) substantial evidence for, 10-100 (1-2) strong evidence for, >100 (>2) decisive evidence for.

The Bayes factor was calculated from the MCMC samples as the Savage-Dickey density ratio^46,47^,

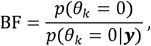

where *p*(*θ_k_* = 0) and *p*(*θ_k_* = 0|***y***) were estimated from the prior and posterior samples using, in each case, a logspline density estimate^48^. Sampling from the posterior is described above; for the prior, we generated 4 × 10^6^ samples from *p*(*θ_k_*) using custom R code.

## Results

### HFD negatively affects metabolic phenotypes in male and female HS rats

As expected, HS males and females on HFD showed negative effects of HFD on multiple metabolic phenotypes relative to rats on LFD **(Fig. 1a-j).** These include increased final body weight (males: F_(1, 323)_ = 48.89, p < 0.0001, females: F_(1, 160)_ = 21.67, p = < 0.0001), increased retroperitoneal fat pad weight (males: F_(1, 296)_ = 122.83, p < 0.0001, females: F_(1, 136)_ = 27.96, p < 0.0001), decreased glucose tolerance (males: F_(1, 233)_ = 30.69, p < 0.0001, females: F_(1, 160)_ = 46.96, p < 0.0001), and increased fasting glucose (males: F_(1, 216)_ = 12.36, p = 0.0007, females: F_(1, 160)_ = 30.86, p < 0.0001), while fasting insulin was only affected in males (males: F_(1, 177)_ = 7.75, p = 0.0064, females: F_(1, 61)_ = 0.024, p = 0.88). Additional fat pads, gonadal and omental, were also significantly increased in HFD males and females (gonadal male: F_(1, 296)_ = 59.16, p < 0.0001, gonadal female: F_(1, 136)_ = 41.80, p < 0.0001, omental male: F_(1, 290)_ = 36.27, p < 0.0001, omental female: F_(1, 136)_ = 11.49, p = 0.0012, data not shown).

**Figure 1.**
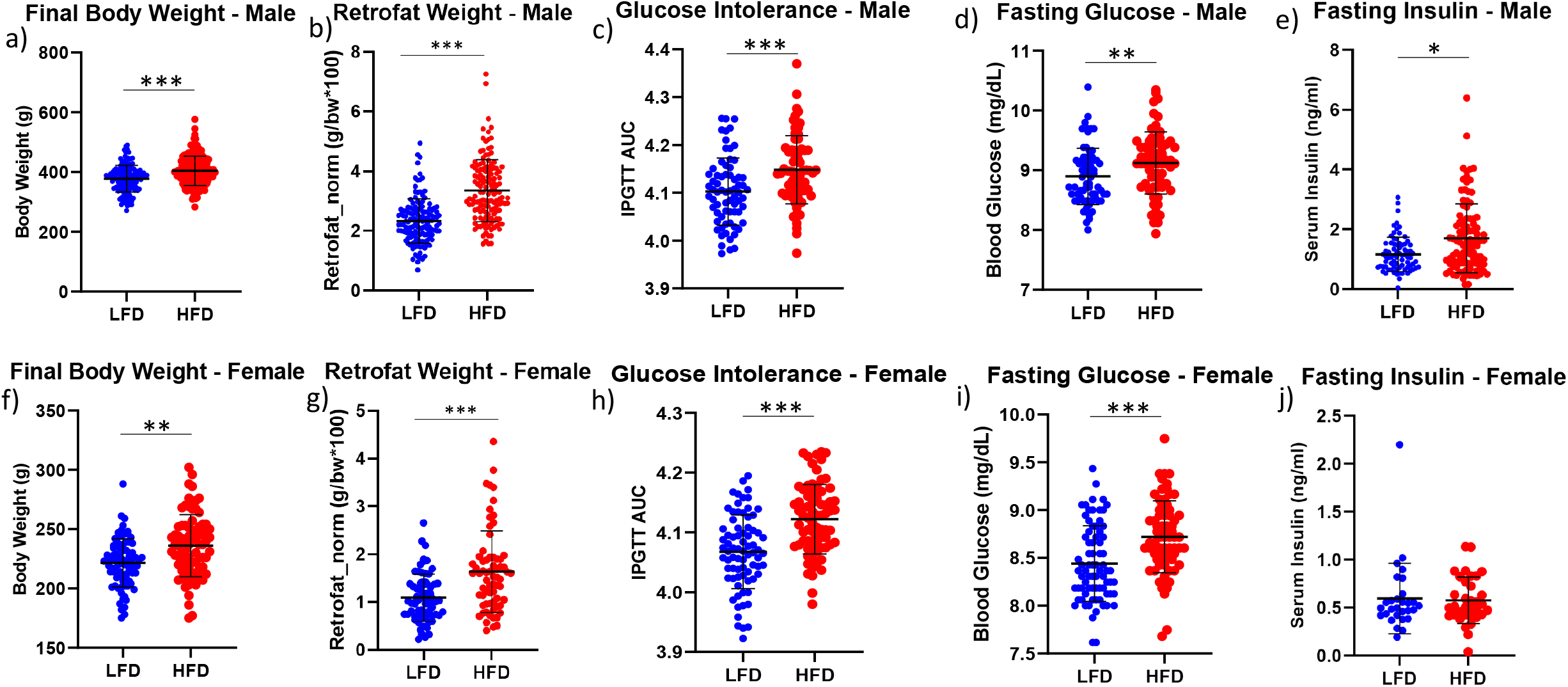
Metabolic effects of high fat diet in male and female HS rats. Final body weight, retroperitoneal fat pad weight (retrofat), intraperitoneal glucose tolerance test AUC (glucose intolerance), fasting glucose, and fasting insulin are shown for (a-e) male and (f-j) female HS rats. Each dot represents an individual animal (blue = LFD, red = HFD). Mean for each group with standard deviation are shown in black. ***< 0.0001; ** <0.001; * <0.05. Sample sizes are provided in supplementary Table 1.

### Effect of HFD on OFT anxiety-like behaviors and FST passive-coping behaviors found only in male HS rats

Anxiety-like behavior was measured via EPM and OFT. Male rats on HFD showed increased anxiety-like behavior in the OFT, while no effect of diet was seen in females **(Fig. 2a-f).** Specifically, male rats on HFD showed significantly decreased rearing episode count (F_(1, 239)_ = 3.91, p = 0.0499) and a trend toward increased rest episode count (F_(1, 239)_ = 3.72, p = 0.056), and movement episode count (F_(1, 239)_ = 3.57, p = 0.061) relative to male rats on LFD. Despite increased rest and movement episode count, male rats on HFD showed a trend toward decreased total distance moved at 5-minutes (F_(1, 239)_ = 2.98, p = 0.086, data not shown) and 30 minutes (F_(1, 239)_ = 3.78, p = 0.055, data not shown). When retroperitoneal fat pad weight was included as a fixed effect co-variate in the model, no trait remained or trended toward significant, indicating that the diet effect is likely acting through increased fat pad size. In contrast to males, diet had no effect on anxiety-like behavior in the OFT in females (rest episode count: F_(1, 184)_ = 0.209, p = 0.65, rearing episode count: F_(1, 184)_ = 0.49, p = 0.49, and movement episode count: F_(1, 184)_ = 0.19, p = 0.66). Although we saw effects of diet on anxiety-like behavior in males in the OFT, there was no effect of diet on EPM closed arm time **(Table 2** males: F_(1, 251)_ = 1.64, p = 0.20, females: F_(1, 180)_ = 0.090, p = 0.77) or open arm time (males: F_(1, 251)_ = 2.08, p = 0.15, females: F_(1, 180)_ = 0.28, p = 0.60) in either males or females.

**Figure 2.**
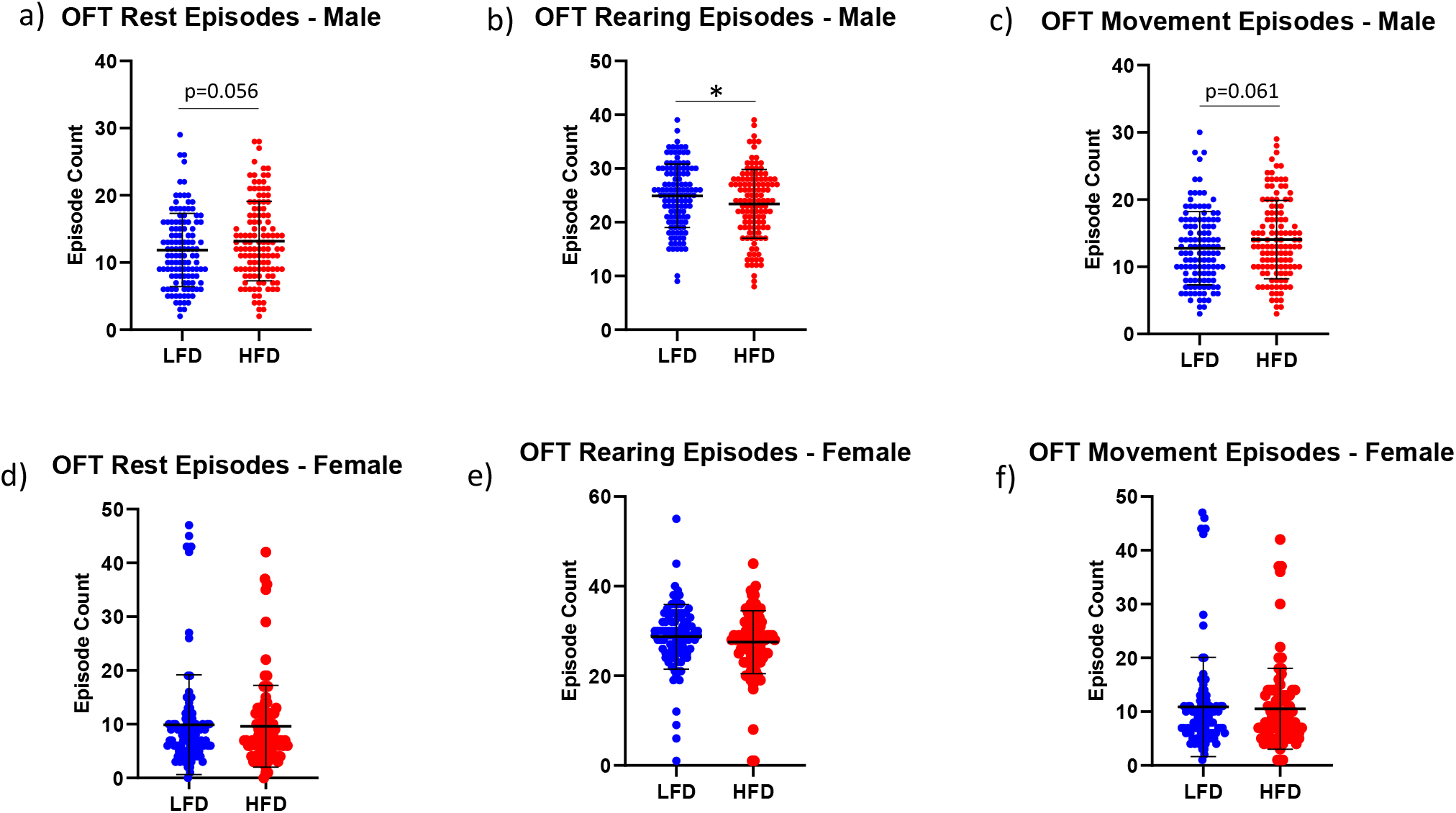
Sex-specific effects of high fat diet on male, but not female, HS rats in the open field test. Anxiety-like behavior was measures via rest episode count, rearing episode count, and movement episode count in the open field test (OFT) for (a-c) male and (d-f) female HS rats following 9 weeks of diet consumption. Each dot represents an individual animal (blue = LFD, red = HFD). Means for each group with standard deviation are shown in black. * <0.05. Sample sizes are provided in supplementary Table 1.

**Table 2.** No effect of high fat diet on elevated plus maze in HS rats.

| Sex | Phenotype | HFD (±SD) | LFD (±SD) |
| --- | --- | --- | --- |
| Male | EPM Open arm time | 85.86 (51.5) | 93.09 (51.0) |
|  | EPM Closed arm time | 208.25 (51.2) | 201.62 (50.5) |
| Female | EPM Open arm time | 99.09 (43.6) | 103.01 (48.3) |
|  | EPM Closed arm time | 196.34 (43.0) | 193.80 (48.1) |

Mean ± standard deviation is shown.

Passive-coping behavior was measured using the FST. HFD significantly affected passive-coping behavior in the FST in males but not females **(Fig. 3a-f).** Specifically, males on HFD showed increased floating (F_(1, 315)_ = 30.81, p < 0.0001), decreased climbing (F_(1, 315)_ = 6.32, p = 0.013), and decreased swimming (F_(1, 315)_ = 25.33, p < 0.0001) relative to males on LFD. In contrast to males, diet had no effect on FST behavior in females (floating: F_(1, 160)_ = 0.53, p = 0.47, climbing: F_(1, 160)_ = 0.91, p = 0.34, and swimming: F_(1, 160)_ = 0.45, p = 0.51). When retroperitoneal fat pad weight was included as a fixed effect in the model, it had a significant effect for FST floating (males: F_(1, 315)_ = 31.88, p < 0.0001, females: F_(1, 160)_ = 15.07, p < 0.0001) and swimming (males: F_(1, 315)_ = 34.64, p < 0.0001, females: F_(1, 160)_ = 20.81, p = 0.0002), in both males and females, but not climbing (males: F_(1, 315)_ = 0.31, p = 0.58, females: F_(1, 160)_ = 0.025, p = 0.88). The effect of diet on floating in males remained significant, but at a less significant p-value (F_(1, 315)_ = 4.15, p = 0.043) indicating that the effect of diet is driven, at least partially, by fat pad weight. Diet was no longer statistically significant in males when retroperitoneal fat was included as a fixed effect for FST swimming (F_(1, 315)_ = 2.21, p = 0.14), although trended toward significant for climbing (F_(1, 315)_ = 3.43, p = 0.065).

**Figure 3.**
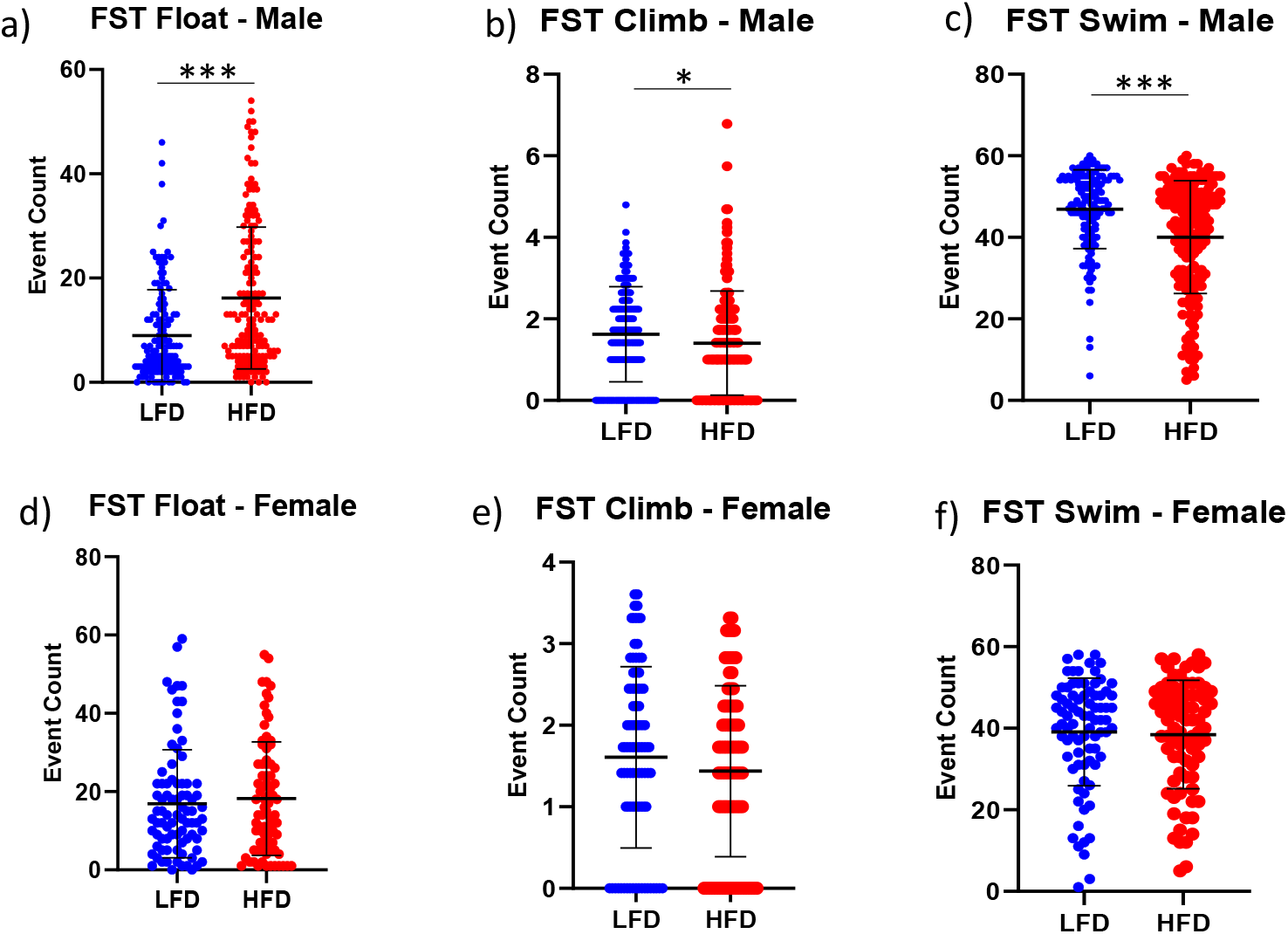
Sex-specific effects of high fat diet on male, but not female, HS rats in the forced swim test. Passive-coping behavior was measured via float, climb, and swim in the forced swim test (FST) for (a-c) male and (d-f) female HS rats following 11 weeks of diet consumption. Each dot represents an individual animals (blue = LFD, red = HFD). Means for each group with standard deviation are shown in black. *** < 0.0001; * <0.05. Sample sizes are provided in supplementary Table 1.

### Factor analysis demonstrates independence between metabolic and behavioral phenotypes and supports sex-dependent effect of diet on behavioral outcomes

To probe the data for underlying latent or hidden factors, behavioral and metabolic phenotypes were analyzed via factor analysis. We found that the metabolic and behavioral phenotypes were mostly not related by any underlying factors in males **(Table 3)** or females **(Table 4).** The loadings for each factor came from metabolic measures or measures of a single behavioral test, except for factor 5 in males **(Table 3,** males: Factor 1 = OFT, Factor 2 = metabolic, Factor 3 = EPM, Factor 4 = FST, Factor 5 = FST/OFT; **Table 4** females: Factor 1 = EPM, Factor 2 = OFT, Factor 3 = metabolic, Factor 4 = FST, and Factor 5 = FST). While limited (male factor 5 explains 9.3% of total variance), the results in males suggest a latent factor relating decreased FST climbing and increased OFT hyperactivity. Despite the strong correlation between FST immobility and retroperitoneal fat shown below, the factor analysis demonstrates that behavioral and metabolic traits are largely independent.

**Table 3.** Factor analysis shows little evidence of underlying relationships between metabolic and behavioral testing in male HS rats.

| Category | Phenotype | Factor1 | Factor2 | Factor3 | Factor4 | Factor5 |
| --- | --- | --- | --- | --- | --- | --- |
| Metabolic | Final body weight | -0.0081 | 0.5556 | -0.1578 | -0.1713 | -0.0192 |
|  | Retroperitoneal fat | -0.1140 | 0.8868 | 0.0632 | 0.1836 | 0.0542 |
|  | Gonadal fat | -0.0932 | 0.8857 | -0.0339 | 0.0720 | -0.0114 |
|  | Omental fat | -0.0264 | 0.9360 | 0.0121 | 0.0024 | 0.0289 |
|  | Fasting insulin | 0.0453 | 0.2351 | -0.0213 | 0.0985 | -0.0986 |
|  | Fasting glucose | -0.0327 | 0.3273 | -0.2066 | 0.1855 | -0.0417 |
|  | IPGTT AUC | 0.0635 | 0.3920 | 0.0866 | 0.1972 | -0.0488 |
| EPM | Closed arm time | 0.0304 | 0.0315 | -0.9832 | -0.0012 | 0.0556 |
|  | Open arm time | -0.0426 | -0.0206 | 0.9836 | 0.0148 | -0.0456 |
|  | Closed arm entries | 0.1972 | 0.0273 | -0.5253 | 0.1970 | 0.0281 |
|  | Open arm entries | 0.0108 | -0.0432 | 0.7684 | 0.0710 | 0.0822 |
| FST | Swim | 0.0740 | -0.1589 | 0.0229 | -0.9626 | 0.0616 |
|  | Climb | -0.0430 | -0.2093 | 0.2114 | -0.1471 | -0.4062 |
|  | Float | -0.0511 | 0.2139 | -0.0739 | 0.9522 | 0.0417 |
| OFT (5min) | Rest episode count (5min) | -0.5435 | -0.0951 | 0.0159 | -0.0913 | 0.8132 |
|  | Movement episode count (5min) | -0.5243 | -0.0843 | 0.0139 | -0.0860 | 0.8268 |
|  | Rearing episode count (5min) | 0.8212 | 0.0020 | -0.0648 | 0.0410 | -0.1729 |
|  | Total distance (5min) | 0.7656 | 0.0567 | -0.1487 | -0.0579 | -0.3417 |
|  | Center time (5min) | -0.3752 | 0.0118 | 0.0438 | 0.0827 | -0.0277 |
| OFT (30min) | Total distance (30min) | 0.8952 | -0.0364 | -0.0420 | 0.0013 | -0.0608 |
|  | Movement episode count (30min) | 0.1786 | -0.1762 | 0.0719 | -0.0428 | 0.5608 |
|  | Rearing episode count (30min) | 0.7985 | -0.0935 | 0.0496 | 0.0447 | 0.1601 |

Loadings shown for metabolic and behavioral measures assessed for latent factors in HS males. Measures most associated with a given factor ( $\pm 0.4$ ) are shown in green (positive association) and red (negative association).

**Table 4.**
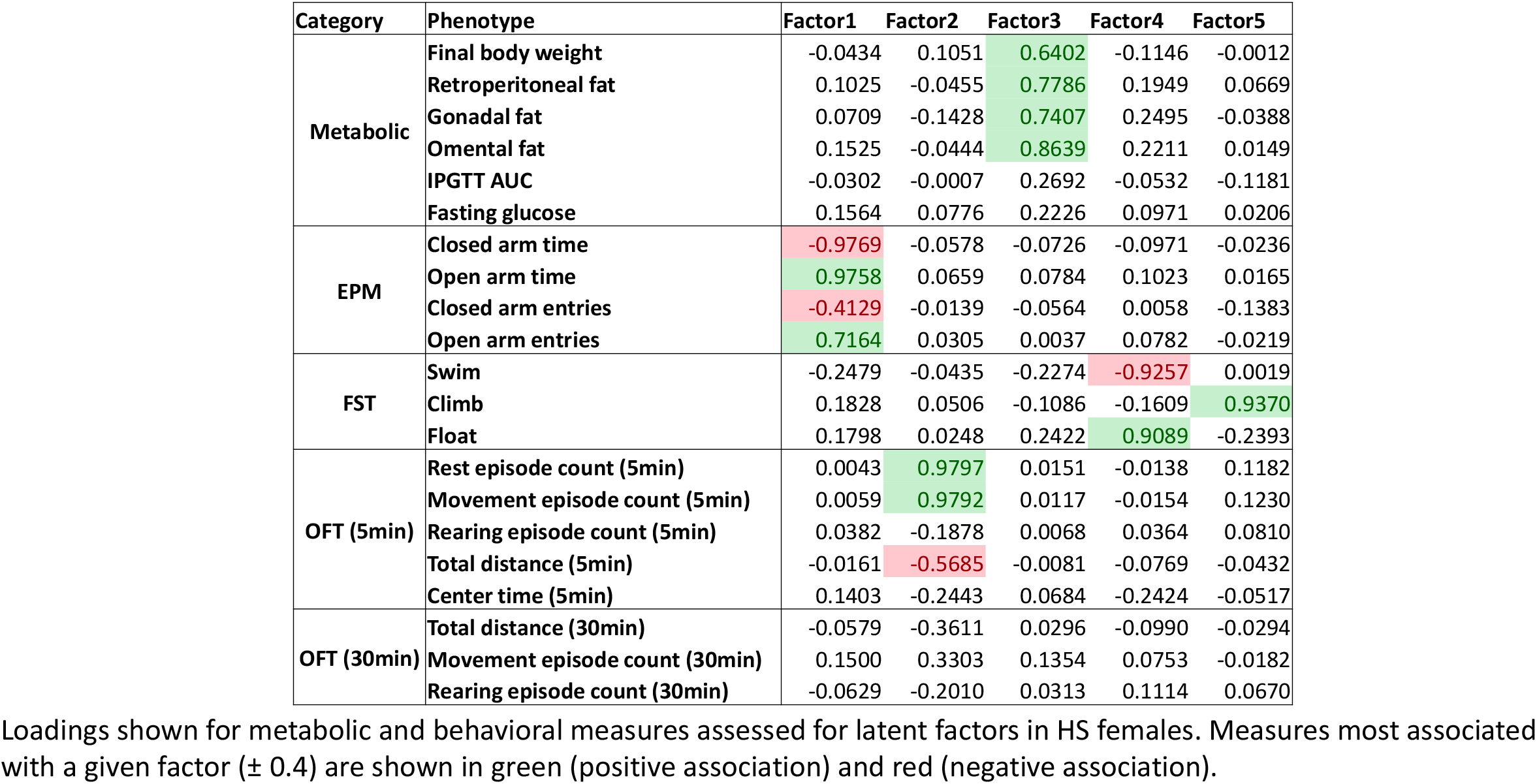
Factor analysis shows no evidence of underlying relationships between metabolic and behavioral testing in female HS rats.

| Category | Phenotype | Factor1 | Factor2 | Factor3 | Factor4 | Factor5 |
| --- | --- | --- | --- | --- | --- | --- |
| Metabolic | Final body weight | -0.0434 | 0.1051 | 0.6402 | -0.1146 | -0.0012 |
|  | Retroperitoneal fat | 0.1025 | -0.0455 | 0.7786 | 0.1949 | 0.0669 |
|  | Gonadal fat | 0.0709 | -0.1428 | 0.7407 | 0.2495 | -0.0388 |
|  | Omental fat | 0.1525 | -0.0444 | 0.8639 | 0.2211 | 0.0149 |
|  | IPGTT AUC | -0.0302 | -0.0007 | 0.2692 | -0.0532 | -0.1181 |
|  | Fasting glucose | 0.1564 | 0.0776 | 0.2226 | 0.0971 | 0.0206 |
| EPM | Closed arm time | -0.9769 | -0.0578 | -0.0726 | -0.0971 | -0.0236 |
|  | Open arm time | 0.9758 | 0.0659 | 0.0784 | 0.1023 | 0.0165 |
|  | Closed arm entries | -0.4129 | -0.0139 | -0.0564 | 0.0058 | -0.1383 |
|  | Open arm entries | 0.7164 | 0.0305 | 0.0037 | 0.0782 | -0.0219 |
| FST | Swim | -0.2479 | -0.0435 | -0.2274 | -0.9257 | 0.0019 |
|  | Climb | 0.1828 | 0.0506 | -0.1086 | -0.1609 | 0.9370 |
|  | Float | 0.1798 | 0.0248 | 0.2422 | 0.9089 | -0.2393 |
| OFT (5min) | Rest episode count (5min) | 0.0043 | 0.9797 | 0.0151 | -0.0138 | 0.1182 |
|  | Movement episode count (5min) | 0.0059 | 0.9792 | 0.0117 | -0.0154 | 0.1230 |
|  | Rearing episode count (5min) | 0.0382 | -0.1878 | 0.0068 | 0.0364 | 0.0810 |
|  | Total distance (5min) | -0.0161 | -0.5685 | -0.0081 | -0.0769 | -0.0432 |
|  | Center time (5min) | 0.1403 | -0.2443 | 0.0684 | -0.2424 | -0.0517 |
| OFT (30min) | Total distance (30min) | -0.0579 | -0.3611 | 0.0296 | -0.0990 | -0.0294 |
|  | Movement episode count (30min) | 0.1500 | 0.3303 | 0.1354 | 0.0753 | -0.0182 |
|  | Rearing episode count (30min) | -0.0629 | -0.2010 | 0.0313 | 0.1114 | 0.0670 |

Loadings shown for metabolic and behavioral measures assessed for latent factors in HS females. Measures most associated with a given factor ( $\pm 0.4$ ) are shown in green (positive association) and red (negative association).

Factor loadings were then used to calculate “factor phenotypes” to analyze the effect of diet on metabolic and behavioral outcomes using fewer dimensions. Consistent with diet effect on individual phenotypes, diet had a significant effect on factors 1 (OFT), 2 (metabolic), and 4 (FST) in males (factor 1 (OFT): F_(1, 323)_ = 5.41, p = 0.022, factor 2 (metabolic): F_(1, 323)_ = 47.72, p < 0.0001, factor 3 (EPM): F_(1, 323)_ = 1.90, p = 0.17, factor 4 (FST): F_(1, 323)_ = 25.34, p < 0.0001, factor 5 (FST/OFT): F_(1, 323)_ = 0.0002, p = 0.99). Notably, the only factor related to multiple tests, male factor 5, was not significantly affected by diet despite most behaviors associated with factor 5 being affected by diet individually. In contrast to males, only factor 3 (metabolic) was identified in females as significantly affected by diet (factor 1 (EPM): F_(1, 186)_ = 0.17, p = 0.68, factor 2 (OFT): F_(1, 186)_ = 0.19, 0.66, factor 3 (metabolic): F_(1, 186)_ = 24.89, p < 0.0001, factor 4 (FST): F_(1, 186)_ = 0.45, p = 0.51, factor 5 (FST): F_(1, 186)_ = 0.91, p = 0.34). These studies support the above findings showing an effect of diet on behavior in males, but not females.

### Correlation analysis

We found strong correlations between metabolic measures and for behavioral measures within the same behavioral test in both male and female HS rats **(Supp. Table 3 and 4).** We also saw strong correlations between fat pad weight and FST behavior in both sexes and across diet conditions. Specifically, there was a positive correlation between retroperitoneal fat pad weight and FST immobility in both HFD males **(Supp. Table 3** r = 0.354, p < 0.0001) and LFD males **(Supp. Table 3** r = 0.385, p < 0.0001). Female HS rats showed a positive correlation between retroperitoneal fat pad weight and FST immobility when both HFD and LFD were analyzed together **(Supp. Table 4** r = 0.325, p = 0.0001), as well as only HFD **(Supp. Table 4** r = 0.370, p = 0.0019), but female LFD did not **(Supp. Table 4** r = 0.255, p = 0.033). Gonadal fat pad weight also significantly correlated with FST immobility for all females, regardless of diet **(Supp. Table 4** r = 0.409, p < 0.0001). Although there were generally few significant correlations across outcomes for separate behavioral tests in males (in support of the factor analysis above), we did find a positive correlation between EPM closed arm entries and OFT total distance in male rats **(Supp. Table 3** r = 0.192, p = 0.0027). Notably, there were no correlations between male FST climbing and OFT measures, in contrast to male factor 5 from the factor analysis above.

Interestingly, although diet had a significant effect on metabolic but not behavioral phenotypes in females, there were significant correlations between adiposity measures and EPM, FST, and OFT measures **(Supp. Table 4).** Additionally, behavioral outcomes were significantly correlated across tests in females. OFT center time negatively correlated with EPM closed arm time **(Supp. Table 4** r = −0.222, p = 0.009). FST swimming in females positively correlated with EPM closed arm time **(Supp. Table 4** r = 0.278, p = 0.0004) and negatively correlated with EPM open arm time (r = −0.288, p = 0.0003), suggesting that FST swimming may represent a marker of anxiety in female rats. Correlation results presented were corrected for multiple comparisons via Benjamini-Hochberg.

### Metabolic and behavioral phenotypes are heritable in both sexes with some gene by diet interactions for behavior in males

Heritability was estimated as “overall heritability” and “heritability of response to diet” for each phenotype. Bayes factors (BF) were calculated for each heritability estimate to measure the evidence for or against the presence of heritability for a phenotype. Heritability for body weight, retroperitoneal fat pad weight, and fasting glucose in male and female rats ranged from 37.0% to 65.3% and was consistent with previous work in this population^24,26^ (see **Table 5;** body weight and retroperitoneal fat pad weight shown in **Fig. 4),** with BF values showing strong evidence for overall heritability of these phenotypes (logBF = 2.45 to 3.54; see **Table 5).** Fasting insulin and IPGTT_AUC in males appeared heritable, but to a lesser extent (20.2% and 23.7%, respectively), with inconclusive support provided by BF values for the presence of heritability (logBF = −0.07 and −0.01, respectively; see **Table 5).** Heritability was not estimated in female fasting insulin due to small sample size. We saw minimal evidence of heritability of response to diet for the metabolic phenotypes in both males and females (1.6% to 12.8%; see **Table 5)** and this was generally supported by BF analysis (logBF = −0.28 to −1.31; see **Table 5).**

**Figure 4.**
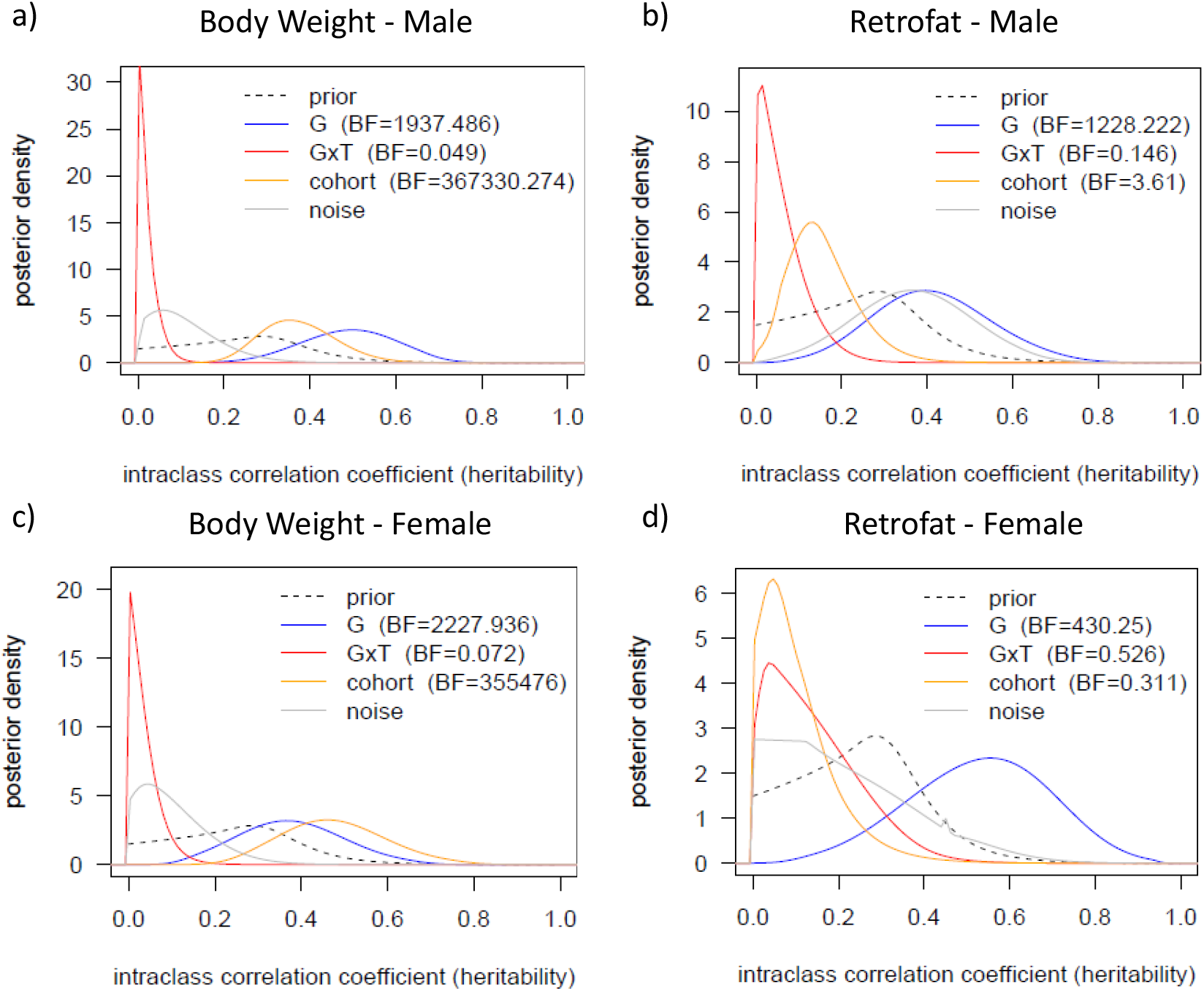
Heritability estimates of body weight and retroperitoneal fat pad weight in male and female HS rats. Posterior distributions for estimates of heritability (G, blue line), gene by diet interaction (GxT, red line), cohort effect (yellow line), and noise (grey line) for retroperitoneal fat (retrofat) and body weight in males (a, b) and females (c, d). The prior distribution is shown via dotted line. Bayes factors (BF) are provided in each figure legend.

**Table 5.** Overall heritability and heritability in response to diet for metabolic and behavioral measures in male and female HS rats.

|  |  |  | Diet | Overall Heritability |  | Gene by Diet Heritability |  |
| --- | --- | --- | --- | --- | --- | --- | --- |
| Sex | Outcome Cat. | Phenotype | p-value | Median (SD) | LogBF | Median (SD) | LogBF |
| Male | Metabolic | Body Weight | 9.154E-11 | 0.690 (0.118) | 3.29 | 0.019 (0.027) | -1.31 |
|  |  | Fasting Glucose | 0.00066 | 0.653 (0.153) | 2.62 | 0.047 (0.058) | -0.85 |
|  |  | Fasting Insulin | 0.00676 | 0.202 (0.14) | -0.07 | 0.091 (0.09) | -0.77 |
|  |  | IPGTT AUC | 1.796E-07 | 0.237 (0.159) | -0.01 | 0.052 (0.061) | -0.79 |
|  |  | Retroperitoneal Fat | <2e-16 | 0.408 (0.137) | 3.09 | 0.055 (0.055) | -0.83 |
|  | Behavior | FST Climb | 0.013 | 0.107 (0.09) | -0.36 | 0.051 (0.062) | -1.03 |
|  |  | FST Float | 9.255E-08 | 0.323 (0.136) | 0.97 | 0.243 (0.117) | 1.22 |
|  |  | FST Swim | 1.11E-06 | 0.242 (0.135) | -0.05 | 0.178 (0.107) | 0.14 |
|  |  | OFT Movement Episode Count | 0.0610 | 0.333 (0.166) | 1.29 | 0.138 (0.109) | -0.28 |
|  |  | OFT Rearing Episode Count | 0.0500 | 0.283 (0.134) | 0.73 | 0.199 (0.128) | -0.13 |
|  |  | OFT Rest Episode Count | 0.0560 | 0.322 (0.165) | 0.71 | 0.134 (0.106) | -0.35 |
| Female | Metabolic | Body Weight | 0.0018 | 0.665 (0.147) | 3.35 | 0.064 (0.065) | -1.14 |
|  |  | Fasting Glucose | 3.629E-07 | 0.54 (0.173) | 2.45 | 0.05 (0.063) | -0.70 |
|  |  | Fasting Insulin | 0.8774 |  |  |  |  |
|  |  | IPGTT AUC | 1.747E-09 | 0.611 (0.152) | 3.54 | 0.073 (0.079) | -0.49 |
|  |  | Retroperitoneal Fat | 1.41E-06 | 0.536 (0.167) | 2.63 | 0.128 (0.105) | -0.28 |
|  | Behavior | FST Climb | 0.344 | 0.199 (0.151) | -0.30 | 0.133 (0.122) | 0.02 |
|  |  | FST Float | 0.467 | 0.656 (0.163) | 2.10 | 0.057 (0.061) | -0.87 |
|  |  | FST Swim | 0.506 | 0.618 (0.156) | 2.31 | 0.047 (0.059) | -0.99 |
|  |  | OFT Movement Episode Count | 0.662 | 0.06 (0.07) | -0.58 | 0.066 (0.075) | -0.72 |
|  |  | OFT Rearing Episode Count | 0.486 | 0.197 (0.145) | -0.32 | 0.197 (0.128) | -0.05 |
|  |  | OFT Rest Episode Count | 0.648 | 0.058 (0.068) | -0.82 | 0.069 (0.077) | -0.81 |

The effect of high fat diet for each phenotype is shown via p-value. Heritability and gene by diet interaction estimates are shown via median ( ± standard deviation, SD). Bayes factors are highlighted for degree of support provided to the respective estimate (green = positive, red = negative, yellow = inconclusive).

When behavioral outcomes were analyzed for overall heritability, we found heritability for OFT measures in males (28.3% – 33.3%) and possibly OFT rearing in females (19.7%), but not in other female OFT measures (5.8% - 6.0%). This was supported by BF analysis **(Table 5).** For FST measures, we found moderate overall heritability of FST swimming and floating in both sexes, with greater heritability in females (61.8% and 65.6%, respectively) relative to males (24.2% and 32.3%, respectively) and this was generally supported by BF analysis. Heritability for FST climbing was lower in both sexes (10.7% in males and 19.9% in females) and BF support was inconclusive.

FST floating in males was the only behavioral outcome to show clear evidence of heritability of response to diet **(Fig. 5b,** 24.3%, logBF = 1.22). OFT measures and FST swimming also exhibited some degree of heritability in response to diet in males (13.4 – 19.9%), although BF analysis remained inconclusive for these measures **(Fig. 5** and **Table 5).** In contrast, with the exception of OFT rearing (19.7%) and FST climbing (13.3%), both with inconclusive BF, there was no evidence for heritability in response to diet for the behavioral measures in females.

**Figure 5.**
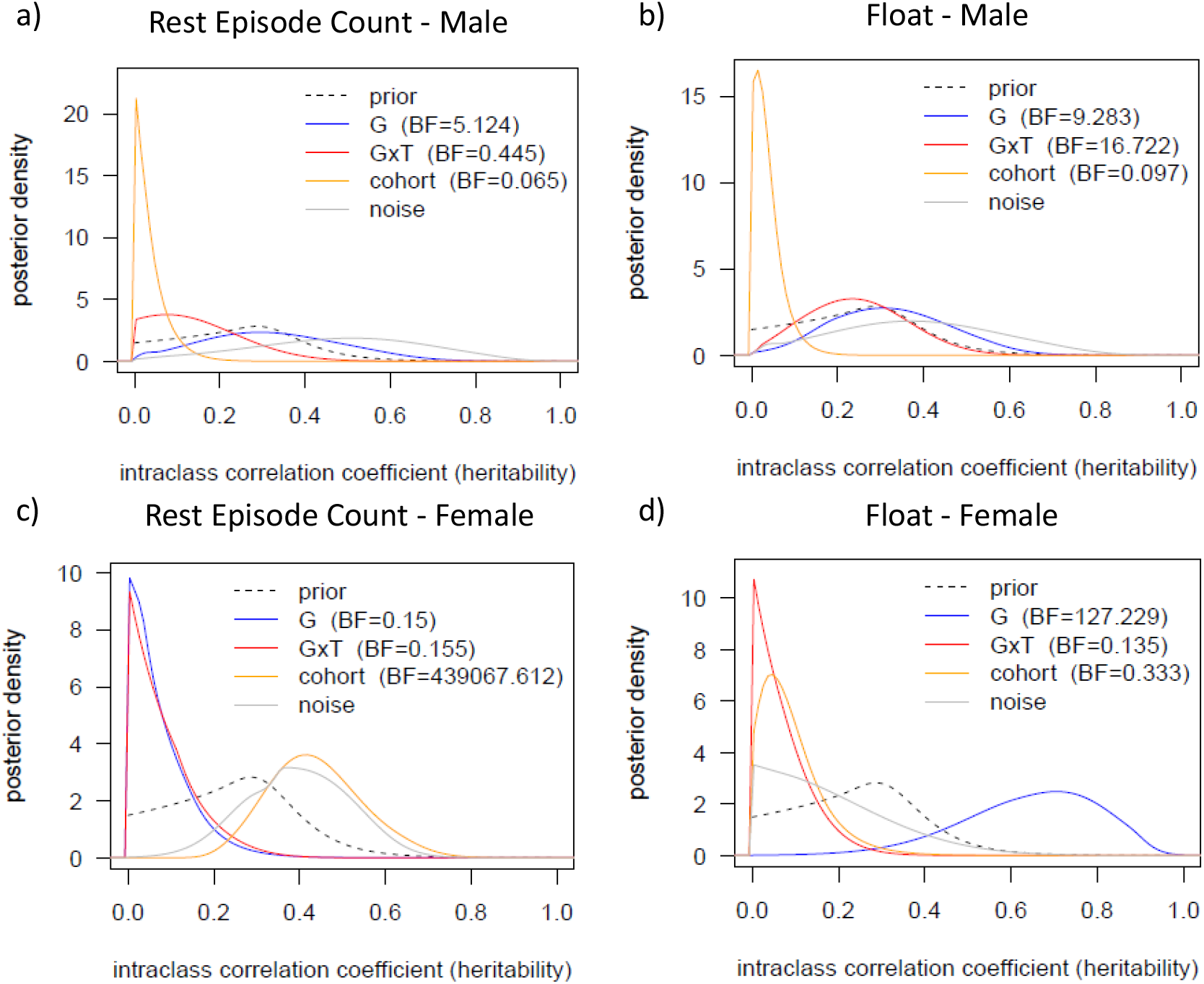
Heritability estimates of FST float and OFT rest episodes in male and female HS rats. Posterior distributions for estimates of heritability (G, blue line), gene by diet interaction (GxT, red line), cohort effect (yellow line), and noise (grey line) for FST float and OFT rest episode count in males (a, b) and females (c, d). The prior distribution is shown via dotted line. Bayes factors (BF) are provided in each figure legend.

## Discussion

In this study, we found that HFD consumption led to worsened metabolic outcomes in both male and female HS rats, while negative behavioral response to HFD was found only in HS male rats. Most metabolic responses showed overall heritability in both sexes. Overall heritability was also found for most behavioral measures, with the exception of OFT traits in females. Surprisingly, although highly genetic, metabolic outcomes were largely independent of gene by diet interactions (e.g., the aggregate effect of polygenes on obesity is similar whether on LFD or HFD). In contrast, we found gene by diet interactions for FST immobility and potentially OFT traits in males, with little to no evidence of gene by diet interactions in females. Overall heritability for both metabolic and behavioral outcomes support the use of HS rats for genetic mapping these traits. The current work demonstrates that gene by diet interactions, at least when defined in terms of interactions between diet and additive polygenes, are likely not influencing diet-induced metabolic changes in HS rats, but may play a role in certain sex-specific behavioral differences.

In the current work, we found that both male and female HS rats on HFD exhibit increased body weight and fat pad weight, reduced glucose tolerance, and increased fasting glucose relative to rats on LFD, with increased fasting insulin seen only in males. This replicates previous work in our lab demonstrating that HS males have worsened metabolic health in response to HFD relative to LFD^14^. Similar to our findings, Sprague-Dawley rats also show negative effects of HFD on metabolic outcomes in both male and female rats^17^. The Diversity Outbred (DO) mouse population also shows worsened metabolic effects in both male and female mice ^49–53^. That being said, sex-dependent differences can be found in the severity or type of physiological effect induced by HFD, and this is often strain dependent^15^. For example, male Wistar rats show increased adiposity and worsened glucose tolerance in response to HFD, whereas females do not^16^; C57/BL6J female mice gained more body weight and fat mass following HFD compared with males^54,55^. The Collaborative Cross (CC), a recombinant inbred mouse population, has also shown sex- and strain-dependent differences in metabolic responses to HFD^56,57^. Male mice showed increased body weight gain on HFD compared to chow than female mice, but fasting glucose was proportionally affected between the sexes^56^. Of the metabolic traits measured in our study, we found that fasting insulin was affected by HFD only in male HS rats and not female. This result has also been shown in C57/BL6 male mice developing insulin resistance in response to HFD, but not females^58^.

In contrast to effects of diet on metabolic outcomes in both male and female HS rats, we saw negative effects of diet on behavior only in males. Specifically, similar to our previous work^14^, we found that HFD increased FST immobility and decreased FST swimming and climbing, demonstrating increased passive coping in male HS rats on HFD. In addition, we saw increased OFT rest episodes, and decreased OFT rearing episodes in male HS rats, indicative of anxious hyperactivity^14^. We were unable to replicate previous findings showing effects of HFD on measures from the EPM. These results are similar to findings in Wistar rats which have shown that exposure to HFD for as few as 7 days is sufficient to induce behavioral changes and these changes are sex-dependent^20^. Specifically, Wistar males on HFD show less active coping in FST and increased moving frequency in OFT^20^, similar to our results in HS males. Male Sprague Dawley rats with free access to a high fat food source in adolescence also developed anxiety-like and passive-coping behavior, an effect not found in females^59^. Inbred C57BL/6 mice also demonstrate sex-specific differences in OFT behavioral responses to HFD. Specifically, C57BL/6 males showed decreased overall movement and females showed increased movement in the OFT following HFD^60^. Similarly, HS males given HFD showed a trend toward decreased total movement distance in the OFT while HS females showed no change. In humans, links have been found between diet and psychological health but sex differences are not apparent^61–63^. The lack of clear sex differences in humans may be a result of the many complex environmental and genetic interactions in behavioral health, thus necessitating very large numbers to see an effect.

Interestingly, we find no evidence that genetic background affects metabolic response to HFD in HS rats. These results are surprising considering that previous work has shown a wide range of metabolic responses to HFD in inbred mouse strains, with some strains showing little to no effect while others are negatively impacted^64,65^. Genetic loci that interact with diet have also been identified in DO mice^51,53^, further suggesting that diet interacts with genetics to impact metabolic traits. Lifespan response to HFD has also been shown to be affected by genetic background in the BxD recombinant inbred mouse population, although the timeline on diet was much longer than the present study^66^. The fact that our findings differ from previous work in rodent models may result from the use of different diets relative to other studies^51,53^, the length of time rats were given diet^66^, the phenotypes measured, and/or the underlying genetic architecture of these traits in response to diet in HS rats relative to other models. Specifically, our assessment of gene by diet effects defines “gene” in terms of a heritability-like polygenic component, that is, the sum of small additive effects of genes across the genome. This aggregate construct will be inefficient at capturing gene by diet interactions if those interactions are limited to a relatively small number of genes -- just as QTL mapping would be ill powered to identify specific genes affecting traits that are highly polygenic. Additionally, the HS rat population, while genetically diverse, does not span the extreme genetic diversity of the DO and CC populations, and may have fewer relevant genes segregating.

Importantly, however, gene-environment interactions have been seen in humans^67–72^, but there has been controversy around which environmental factors play a role. In fact, Tyrell et al^73^ recently identified multiple environmental factors that interacted with obesogenic alleles to affect susceptibility to obesity in humans, but, similar to our findings, “Western” diet was not found to be a significant factor, bringing into question these earlier findings^73^. Thus, the importance of gene by diet interactions on metabolic outcomes in humans remains an open question.

The lack of gene by environment interactions in the HS rat suggests that factors other than genetics may be influencing the phenotypic response to HFD. One possible explanation is that HFD is acting epigenetically (as opposed to genetically) by altering methylation patterns on DNA resulting in differently expressed genes to effect metabolic outcomes. Previous work has shown that acute HFD feeding can alter methylation patterns in adipose tissue in humans^74^ and this has been replicated in animal models^75–78^. Previous work also suggests that diet can cause epigenetic alterations which can exacerbate HFD effects on obesity in offspring, even identifying a stronger effect in female mice^79^. Parent-of-origin (PO) effects may also be playing a role as previous work has shown interactions between PO and perinatal diet in CC recombinant inbred intercross lines^80^. Another explanation may be influences of the microbiome, which is altered by HFD^81^ and can influence susceptibility to diet-induced weight gain independent of genetics^82^. The hypotheses that epigenetics (via methylation or PO effect), the microbiome, or some other factor may be modulating effect of diet on metabolic outcomes in HS rats can be explored in future studies.

In contrast to the metabolic phenotypes, we found that FST floating in males showed evidence of gene by diet interactions. Additionally, some gene by diet interaction may be occurring in male FST swimming and the OFT traits. That is, in HS male rats, genetic background appears to influence how diet impacts passive-coping and potentially anxiety-like behaviors. Thus, altering diet may affect these behavioral outcomes in individual rats based on genetic factors. In contrast, in female rats, other than inconclusive evidence for OFT rearing, most of the behavioral traits did not show gene by diet interactions. To our knowledge, this is the first study to report gene by diet interactions for behavioral traits in male rats. In humans, the idea that dietary change may be a viable tool in addressing affective disorders depending on an individual’s genetic background is supported by some, but not all studies^83^.

Although fat pad weight confounds with diet-induced changes in behavioral outcomes, factor analysis results show that metabolic and behavioral phenotypes are largely independent from each other. Furthermore, measures within each behavioral test are largely independent from measures in other behavioral tests, suggesting that there are not shared factors underlying the behavioral phenotypes in the different tests. Previously, we had shown correlations in male HS rats between OFT and EPM behaviors only in animals on HFD^14^. We were unable to replicate that finding in the current study, showing some correlations between OFT and EPM measures, only when animals on both LFD and HFD were analyzed together. One explanation for this difference may be sample size, which has been greatly increased in the results shown here.

In conclusion, we have shown that both male and female HS rats are susceptible to negative metabolic effects of HFD, but only HS males exhibit negative behavioral phenotypes in response to HFD. In addition, while we see no effect of genetic background on diet-induced metabolic outcomes in either sex, behavioral outcomes show gene by diet interactions in males, particularly for FST floating, with limited evidence for gene by diet interactions in females. Future studies could explore the genes potentially connecting behavior in response to diet in HS rats, particularly in male passive-coping behavior. Overall, this work shows important effects of sex, diet, and genetic background on metabolism and behavior in HS rats. This work also adds to the growing literature characterizing sex differences in metabolic and behavioral phenotypes in rodents in response to HFD.

## Data Availability

The data and code to support the results reported here are available on GitHub (https://github.com/valdarlab/h2GxD-hsrats). These data include: phenotype data (by sex) and pedigree information (by sex) for all study mice, and transformations to use for each phenotype (based on Box Cox analysis). All scripts to transform the raw data, produce additive relationship matrices from the pedigrees, run the statistical models, and process results, and a Docker container in which to run the analyses is available in the GitHub repo.

## Acknowledgements

We acknowledge financial support from the National Institutes of Health (NIH) grant P50 DA037844, which supported the HS rat breeding colony, R35GM127000 supporting ELR and WV, T32GM135123 supporting ELR, and Wake Forest School of Medicine start-up funds which supported the experimental work. We also thank Jeff Weiner’s lab in the Wake Forest Department of Physiology and Pharmacology for the use of the behavioral assays.

The authors declare no competing financial interests.

## Appendix

The Gibbs sampler for fitting Eq.1 is described as follows. For *n* subjects, *p* fixed effect covariates, and *q* random effect covariates (residual error considered separately), the rewrite Eq 1 as:

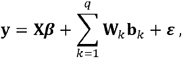

where **X** is an *n × p* matrix of covariate values, ***β*** is a *p* × 1 vector of fixed effects, **W***_k_* = *n×n* diagonal matrix where ***W**_k_* = diag(**d**) when *k* = *G × D* or **W***_k_* = **I** otherwise,

**b***_k_* is an *n* × 1 vector of random effects corresponding to variance component *k*, where 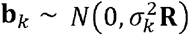 with **R** = **A** for heritability-based components or **R** = **C** for cohort. Compute singular value decomposition **R** = **UDV^T^** and let **M** = **UD**^1/2^, so that we can write **b***_k_* = ***Mα**_k_*, and 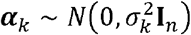. Using prior distributions,

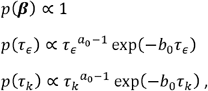

where 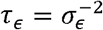 and 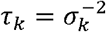, the full conditional distributions for the Gibbs sampler are

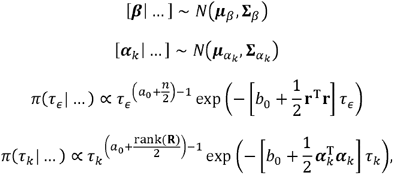

where

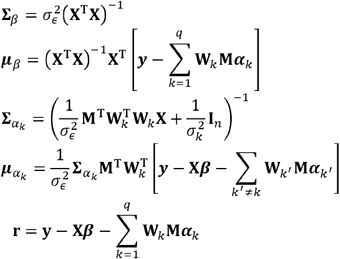

---

**Supplementary Table 1. Sample size for metabolic and behavioral measures.**

**Supplementary Figure 1. Timelines for metabolic and behavioral pilot studies.**

**Supplementary Table 2. Transformation applied to each phenotype prior to analysis.**

**Supplementary Table 3. Correlations between metabolic and behavioral measures for male HS rats.** Metabolic and behavioral measures were assessed for relationships via Pearson correlation in all male data (top), male HFD (middle), and male LFD (bottom). Significantly positive and negative correlations are shown in green and red, respectively. Results are corrected for multiple comparisons via Benjamini-Hochberg procedure (FDR = 0.05).

**Supplementary Table 4. Correlations between metabolic and behavioral measures for female HS rats.** Metabolic and behavioral measures were assessed for relationships via Pearson correlation in all female data (top), female HFD (middle), and female LFD (bottom). Significantly positive and negative correlations are shown in green and red, respectively. Results are corrected for multiple comparisons via Benjamini-Hochberg procedure (FDR = 0.05).

