## Supplemental Figure 1 for "Sex and genetic specific effects on behavioral, but not metabolic, responses to a high fat diet in heterogeneous stock rats"

### Slide 1
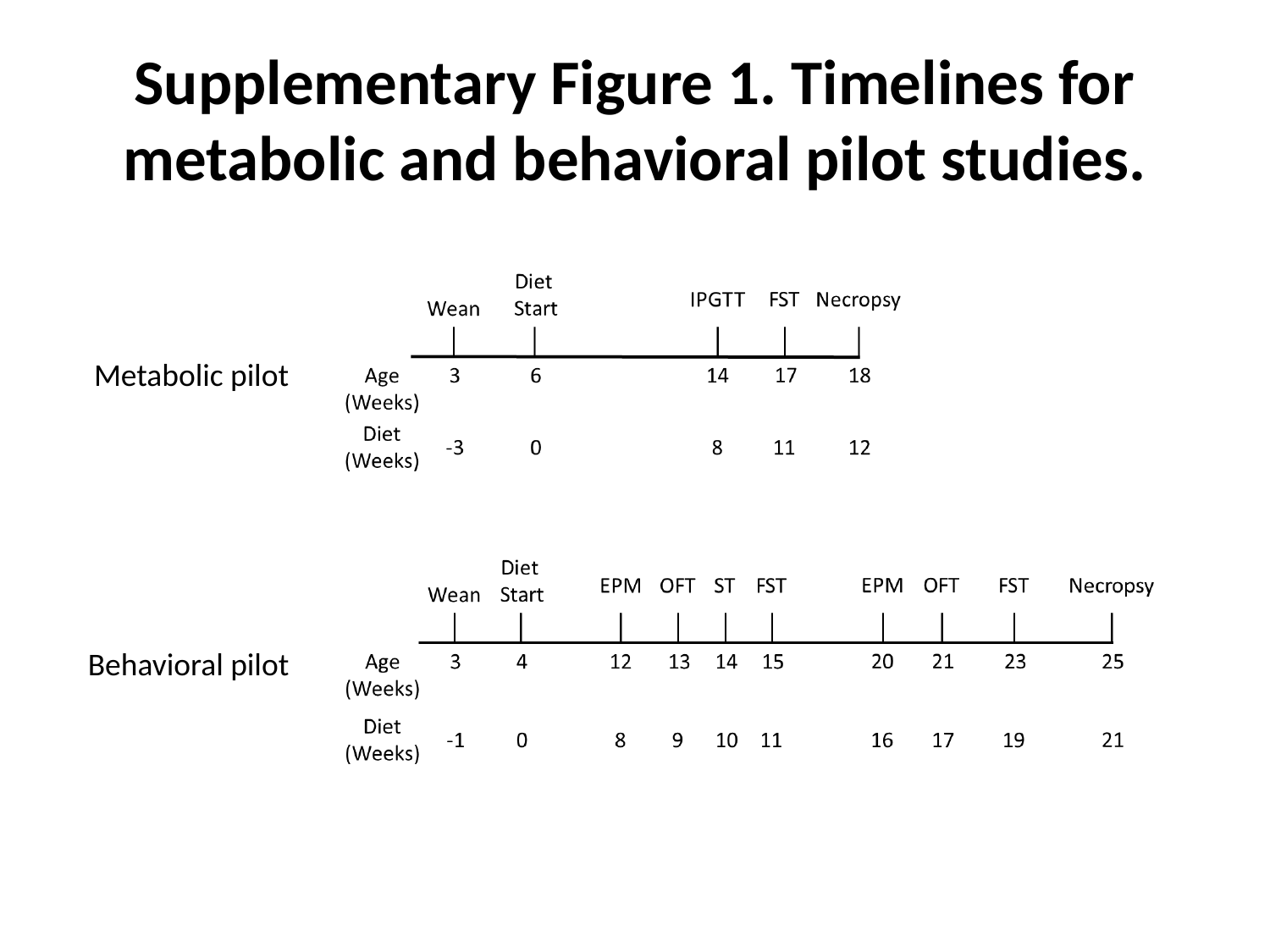

#### Supplementary Figure 1. Timelines for metabolic and behavioral pilot studies.
Metabolic pilot
Behavioral pilot
